# The Fatty Acid Methyl Ester (FAME) profile of *Phytophthora agathidicida* and its potential use as diagnostic tool

**DOI:** 10.1101/2021.04.06.437983

**Authors:** Randy F. Lacey, Blake A. Sullivan-Hill, Julie R. Deslippe, Robert A. Keyzers, Monica L. Gerth

**Author notes:** **Corresponding author**: Address: School of Biological Sciences, Level 2, Te Toki a Rata Building, Victoria University of Wellington, Wellington, 6012.

## Abstract

*Phytophthora* diseases cause devastation to crops and native ecosystems worldwide. In New Zealand, *Phytophthora agathidicida* is threatening the survival of kauri, an endemic, culturally and ecologically important tree species. The current method for detecting *P. agathidicida* is a soil bating assay that is time-consuming and requires high levels of expertise to assess, thus limiting the analytical sample throughput. Here, we characterized the fatty acid methyl ester (FAME) profile of *P. agathidicida*. We also compared it with the FAME profile of *P. cinnamomi* and assessed the efficacy of FAME analysis as a diagnostic tool for detecting the pathogen in soil samples. In FAME analysis, the total fatty acid content is isolated from a sample and converted to FAMEs for analysis, a process that takes less than a day. Unique fatty acid acyl chains can serve as biomarkers for specific organisms. We detected 12 fatty acids in *P. agathidicida*, two of which (20:4ω6 and 20:5ω3) show promise as potential *Phytophthora* specific biomarkers. Collectively, these findings advance our fundamental understanding of *P. agathidicida* biology and provide a promising technique to increase the rate of sample processing and the speed of pathogen detection for *P. agathidicida* in soil.

## Introduction

*Phytophthora agathidicida* is a recently identified plant pathogen that is threatening New Zealand’s native kauri trees (*Agathis australis*) (Weir *et al*., 2015, Bellgard *et al*., 2016). It is the causative agent of kauri dieback disease, which has spread throughout most regions within the natural range of kauri (Bradshaw *et al*., 2020). Kauri trees are massive and can live for thousands of years. Because of this, there can be a latency period from infection to the expression of symptoms (Bradshaw *et al*., 2020). Thus, detection of *P. agathidicida* in soil, before the onset of disease symptoms, is critical for managing the spread of kauri dieback. Currently, there are limited tools available to control the spread of disease, and improved surveillance and diagnostics is an urgent priority for research (Bradshaw *et al*., 2020).

*Phytophthora* can spread in a variety of ways, including by water, root to root contact, and movement of contaminated plant tissue or soil (Erwin & Ribeiro, 1996). The lifecycle of *Phytophthora* species is commonly characterized by the production of vegetative mycelial growth and various spore types (Erwin & Ribeiro, 1996). Zoospores are short-lived, motile spores that initiate infection in roots (Bradshaw *et al*., 2020). Oospores are thick-walled, dormant spores that can survive in soil for years (Bradshaw *et al*., 2020). Each lifecycle stage plays a critical role in disease spread. In New Zealand there are at least 30 known species of *Phytophthora* that cause disease in agricultural, horticultural, and indigenous forest settings (Scott & Williams, 2014). For example, *P. cinnamomi* often co-occurs with *P. agathidicida* at sites of infected kauri (Waipara *et al*., 2013).

Currently the primary method used to detect *P. agathidicida* in soil is a baiting assay (Beever *et al*., 2010). Soil-baiting is a method that is used for detection and isolation of many different *Phytophthora* species (Erwin & Ribeiro, 1996, O’Brien *et al*., 2009). This method is effective but has limitations; it is slow (2-3 weeks) and requires a high level of expertise to correctly identify the colony and spore morphology that distinguishes *P. agathidicida* from other *Phytophthora* species (Beever *et al*., 2010, Bradshaw *et al*., 2020). These limitations reduce sample throughput, which limits the capacity to monitor the geographic spread of *P. agathidicida* in a timely manner. More rapid DNA-based molecular diagnostic tools are in development, but so far have limited effectiveness for detecting *P. agathidicida* in soil samples (Than *et al*., 2013, McDougal *et al*., 2014, Winkworth *et al*., 2020).

Fatty acid methyl ester (FAME) analysis is potentially an alternative or complementary tool for the detection of organisms in soil. FAME analysis has been used extensively to characterize microbial community structure in soils and has also been applied as a diagnostic tool for the detection of specific organisms in environmental samples (Cavigelli *et al*., 1995, Drenovsky *et al*., 2004, Rastogi & Rajesh, 2011, Yousef *et al*., 2012). In FAME analysis, total lipids are extracted from an organism or environmental sample, acyl chains are released as fatty acids and converted to their corresponding methyl esters with subsequent gas chromatography-mass spectrometry (GC-MS) analysis (Welch, 1991, Drenovsky *et al*., 2004). Organisms differ in the types and quantities of acyl chains that are components of their lipids (White *et al*., 2002, Ehrhardt *et al*., 2010). Consequently, the presence of or varying ratios of specific FAMEs can indicate the presence and quantitative abundance of particular taxa in a sample, in which case the FAME or ratio of FAMEs may be considered a biomarker. For example, the neutral lipid 16:1ω5 is widely used as an indicator of arbuscular mycorrhizal fungi in soils (Olsson, 1999). Specific to *Phytophthora*, it was shown that soils with *P. sojae* zoospores added showed increased levels of 18:2ω6, 20:4ω6, and 22:1ω6, indicating that increased ratios of these fatty acids to background fatty acids may potentially serve as a biomarker for the presence of *P. sojae* in soil (Yousef *et al*., 2012).

Additionally, A significant advantage of a FAME approach, relative to soil baiting and PCR-based methods, is the potential to quantify the biomass of the pathogen in the sample. Finally, the characterisation of FAMEs is relatively rapid; the process from a sample to quantitative detection of a biomarker can take less than a day.

Here, we present the results of FAME analysis of *P. agathidicida* across several key lifecycle stages, including mycelia, oospores, and zoospores. Additionally, a comparison of FAME profiles of *P. agathidicida* and *P. cinnamomi* reveals that many fatty acids are conserved between the two species; no fatty acids unique to *P. agathidicida* were identified. However, elevated ratios of the long-chain polyunsaturated fatty acids 20:5ω3 and 20:4ω6 were observed in soil samples with *P. agathidicida* oospores added. This suggests that FAME analysis may be a useful diagnostic tool for the detection of *Phytophthora* species in soil.

## Materials and Methods

### Culture conditions and mycelia production

*P. agathidicida* NZFS 3770 and *P. cinnamomi* NZFS 3910 (obtained from Scion, Rotorua, NZ) were maintained regularly on 10% clarified V8 (cV8) agar plates in the dark at 22 °C. Mycelia is a common tissue source of lipids for FAME profiling of *Phytophthora* species (Larkin & Groves, 2003, Duan *et al*., 2013). Soil microbes may adjust the relative concentrations of membrane lipids to acclimate to different growth temperature (Griffiths *et al*., 2003, Yousef *et al*., 2012). Therefore, we initially characterized the basic fatty acid profile of *P. agathidicida* mycelia at two different temperatures. We selected 16 °C as a value similar to mean annual temperature in the host’s range and 22 °C as *P. agathidicida* shows optimal growth *in vitro* culture at this value. Mycelial mats for lipid extraction were grown in liquid 10% cV8 broth for 48 h at 16 °C and 22 °C in the dark. Mycelia were separated from the agar plug from which growth was initiated and washed in deionized water to remove residual cV8 broth.

### Oospore production

*P. agathidicida* oospores were produced as described in Fairhurst et al. (Fairhurst *et al*., 2021). Briefly, three 3 mm agar plugs were taken from the leading edge of mycelial growth on agar plates and inoculated in 15 mL of 4% w/v carrot broth containing 12 µg/mL β-sitosterol and grown in the dark at 22 °C for two weeks. The resulting mycelial mats were harvested and oospores were isolated by homogenization using a tissue homogenizer for 2 min followed by sonication for 1 min on ice. The homogenized mixture was sequentially filtered through 100 µm and 40 µm filters to separate the oospores from mycelial fragments. The concentration of the filtered oospore suspension (oospores/mL) was estimated using a disposable hemocytometer by averaging three separate counts.

### Zoospore production

*P. agathidicida* zoospores were produced as described in Lacey et al. (Lacey *et al*., 2021). Briefly, mycelial mats were initially grown in 2% w/v carrot broth supplemented with 15 µg/mL β-sitosterol for 30 h. The mycelial mats were then washed with 2% w/v soil solution and incubated under light for 14 h. The soil solution was removed and the mycelial mats were washed with water. Zoospore release was induced by adding ice-cold water and incubating the mycelial mats at 4 °C for 20 min. After sufficient zoospore release, the concentration of spores was estimated using disposable hemocytometers by averaging three separate spore counts.

### Fatty acid extraction, conversion to FAMEs, and GC-MS analysis

For FAME analyses, mycelia (∼200 mg) from *P. agathidicida* and *P. cinnamomi* were grown and prepared as described above in biological triplicate. For *P. agathidicida* oospore and zoospore analyses, each spore type was produced as described above. A total of 250,000 zoospores or 100,000 oospores in 250 µL was used for lipid extraction. For the detection of oospores in soil, we collected five soil samples from forest and garden locations in Wellington, NZ, a region outside the native range of kauri and presumably free of *P. agathidicida*. Soil samples were collected with a trowel, which was sterilized with 70% ethanol between samples. Soil samples were sealed in plastic bags and placed on ice for transport to the laboratory. Samples were then freeze-dried and stored at −20 °C. Laboratory grown oospores were added in 250 µL aliquots to 0.5 g sub-samples of soil sample A. The mixtures of soil and laboratory-grown oospores were subsequently subjected to lipid extraction and FAME production.

For all conditions, lipid extraction and conversion to FAMEs was carried out as described in Duan et al. (2013) with slight modifications. Initially, fatty acids were released and saponified by addition of 3.75 M NaOH dissolved in 50% MeOH_*(aq)*_ (1 mL) to the sample and incubation at 80 °C for 30 min with occasional mixing. Next, free fatty acids were methylated by addition of 62.5% 6 N HCl: 37.5% MeOH solution (2 mL) and incubation at 80 °C for 10 min. Phase separation was then carried out by the addition of 1 mL of a 1:1 mixture of *n*-hexane and methyl-tert butyl ether with vigorous mixing by vortex for 30 sec. After sufficient phase separation, the top organic phase was transferred to a new glass tube and washed with 3 mL of 0.3 M NaOH_(*aq*)_. The top organic phase was again transferred to a new tube and concentrated under a stream of nitrogen gas. The remaining FAMEs were then resuspended in 150 µL *n*-hexane and transferred to 2 mL GC vials with inserts and processed immediately or stored at −20 °C until analysis.

Samples were analysed on a Shimadzu GCMS-QP2010 Plus. The GC column used was a Restek RXI-5SilMS (30 m x 0.25 mm x 0.25 µm) and He was the carrier gas. An aliquot (2 µL) of the sample was injected (8:1 split ratio) at 260 °C with a column flow rate of 1.09 mL/min and linear velocity of 39.6 cm/sec. The GC conditions were as follows: 2 min hold at 150 °C followed by a 10 °C increase per min to 280 °C with a final 10 min hold at 280 °C. The MS transfer line temp was 260 °C. MS analysis began after 4 min and ended at 25 min with 3 scans/sec. Ions between 40 and 600 m/z were detected. The EI ion source operated at 70 eV ionisation energy. Chromatograms were analysed using Shimadzu GCMSsolution Software. FAMEs were annotated using mass spectral matching to the National Institute for Standards and Technology (NIST) 2011 database. These annotations were confirmed based on a comparison of the retention time and fragmentation pattern with those of authentic FAME standards (Larodan). Fatty acids are named using standard nomenclature (fatty acid chain length: number of double bonds followed by an ω, finally the carbon number from the methyl terminus where the first double bond starts, *e*.*g*. linoleic acid ((9*Z*,12*Z*)-octadeca-9,12-dienoic acid) is 18:2ω6).

FAMEs were quantified in two ways. First, the relative amount of each *P. agathidicida* FAME was determined as a percent of the total fatty acid content. To do this, the percent of each FAME was determined by integrating the area under the peak representing each FAME on the chromatogram. The values were then converted to percentages with all identified peaks per sample combined to total 100%. Thus, each FAME is calculated as a fraction of 100% of the total FAMEs identified per sample. Second, we quantified molar amounts of 20:4ω6 and 20:5ω3 in soil samples using a 19:0 fatty acid standard as the internal standard. These two fatty acids were chosen for quantification due to the difference in ratio of 20:5ω3 to 20:4ω6 when comparing soil samples and cultured *P. agathidicida*. For use as an internal standard, 100 nmol of 19:0 fatty acid was added directly to soil samples prior to lipid extraction and conversion to FAMEs. For quantification, response factors for 20:4ω6, 20:5ω3, and 19:0 were first determined and relative response factors between 20:4ω6/19:0 and 20:5ω3/19:0 were subsequently determined (Dodds *et al*., 2005). For each condition, five biological replicates were performed, where each replicate represents a separate extraction, analysed on a separate day. Statistical analyses and graphical representations of the data were performed and generated using Prism GraphPad (Version 8.2.1).

## Results

### Comparison of FAME profiles of P. agathidicida *and* P. cinnamomi

Twelve fatty acids were regularly detected from *P. agathidicida*. The five most abundant fatty acids were 14:0, 16:0, 18:2ω6, 18:1ω9, and 20:5ω3, constituting greater than 75% of the total fatty acid content (Fig. 1). Comparing *P. agathidicida* mycelia grown at 16 °C and 22 °C revealed slight changes in relative fatty acid amounts. Of the five most abundant fatty acids, 14:0, 16:0, and 18:2ω6 were slightly reduced (<5 %) at 16 °C compared to 22 °C while 18:1ω9 and 20:5ω3 were slightly increased (<5 %) at 16 °C compared to 22 °C. We also examined the FAME profile of *P. cinnamomi* mycelia at 16 °C and 22 °C (Suppl. Fig. 1). The FAME profiles of *P. agathidicida* and *P. cinnamomi* were largely similar. Eleven fatty acids were detected in *P. cinnamomi*, all of which overlapped with fatty acids present in *P. agathidicida* (Fig. 1 and Suppl. Fig. 1). Interestingly, the fatty acid 22:1ω9 was regularly present in *P. agathidicida* but not detectable from *P. cinnamomi*. However, previous studies have shown that this fatty acid is present in *P. cinnamomi* at low relative percentages (Duan *et al*., 2013). In general, our fatty acid profile for *P. cinnamomi* was similar to a previously published report that spanned multiple isolates of this species (Duan *et al*., 2013). Duan et al. detected fifteen fatty acids from *P. cinnamomi*. The four additional fatty acids detected in that study comprised less than 2% of the total fatty acid content.

**Figure 1:**
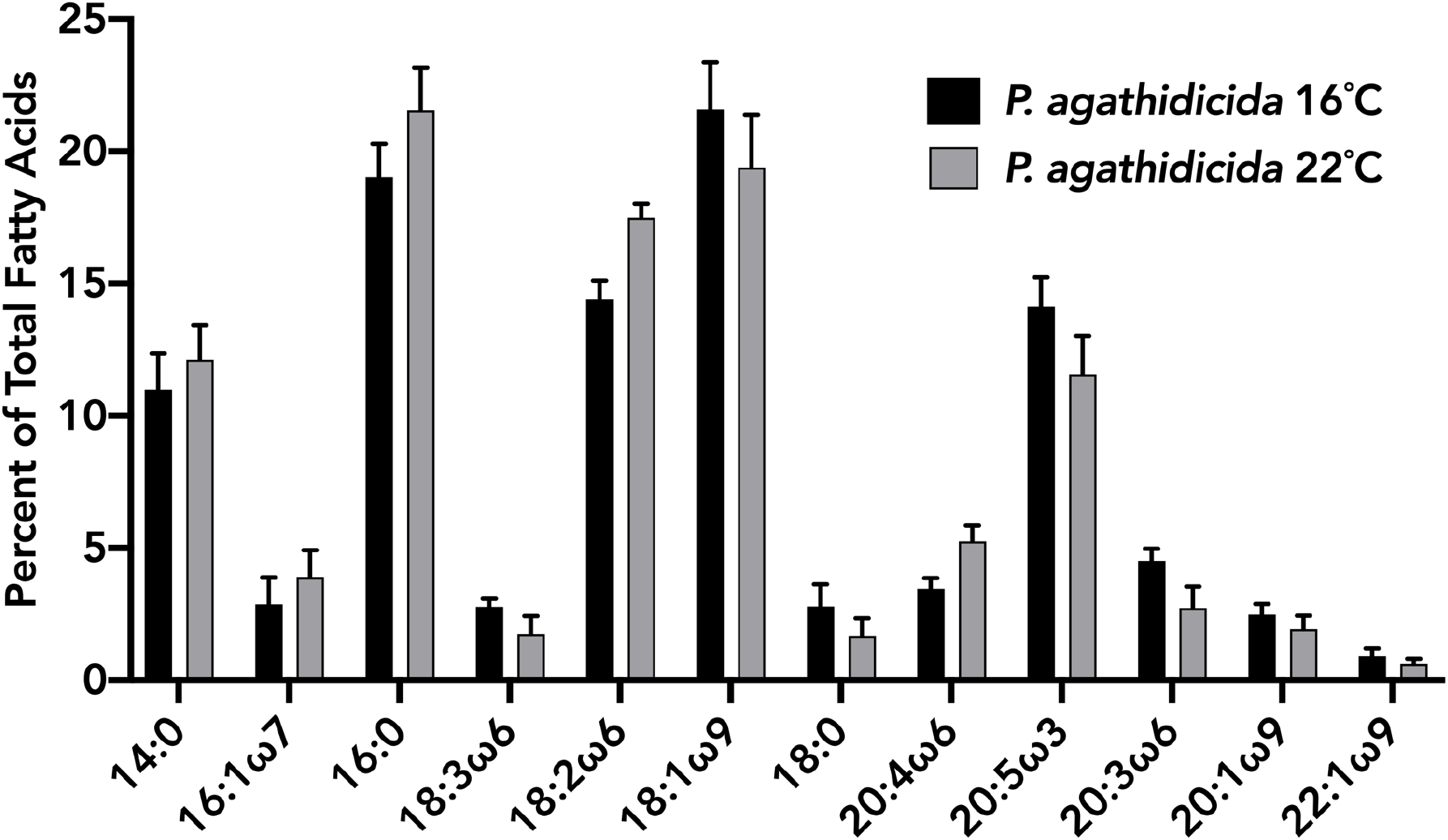
Fatty acid profile of *P. agathidicida* at varying temperatures. FAMEs were produced and analysed from mycelia of *P. agathidicida* grown at 16 °C and 22 °C. The percent of each fatty acid is the average relative percent of five biological replicates. Error bars indicate standard deviation.

### Comparison of FAME profiles of various P. agathidicida *lifecycle stages*

Environmental conditions can affect the relative abundances of oomycete lifecycle stages in soil (Erwin & Ribeiro, 1996). We therefore compared the FAME profiles of *P. agathidicida* oospores, zoospores, and mycelia (Fig. 2). Compared to mycelia, the FAME profiles of oospores and zoospores were less diverse in quality and quantity of acyl chains that were present. Each FAME that was detected from the mycelial samples was also detected in oospores, however, the long-chain unsaturated fatty acids were in lower quantities relative to shorter chain length fatty acids when compared with mycelia. In mycelia, three polyunsaturated, long-chain fatty acids (20:4ω6, 20:5ω3, 20:3ω6) and two mono-unsaturated, long-chain fatty acids (20:1ω9, 22:1ω9) were detected and all except 22:1ω9 constituted >1% of the total FAME profile. In oospores and zoospores, 22:1ω9 was not detected. In zoospores, 18:3ω6 and 20:1ω9 were also not detected. Overall, oospores and zoospores produced lower quantities of long-chain fatty acids. Different growth media are known to induce changes in the FAME profiles of *Phytophthora* species (Larkin & Groves, 2003, Duan *et al*., 2011), and may have contributed to the variation we observed among life-cycles stages. However, these effects are difficult to quantify since no single laboratory procedure exists to produce oospore and zoospores in *P. agathidicida*.

**Figure 2:**
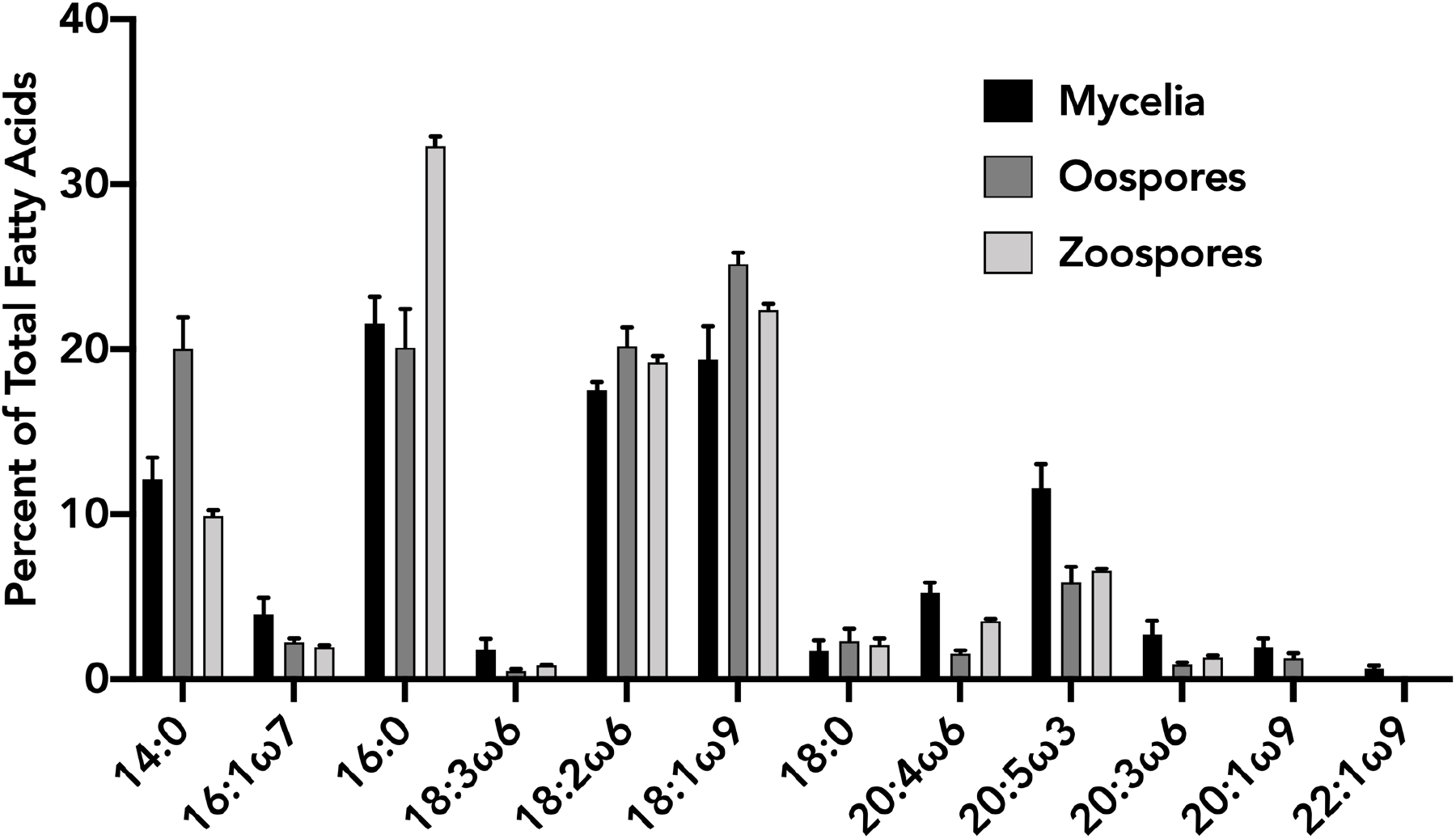
Fatty acid profile comparison of different *P. agathidicida* lifecycle stages. FAMEs were produced and analysed from *P. agathidicida* mycelia, oospores, and zoospores. The percent of each fatty acid is an average relative percent of five biological replicates. Error bars indicate standard deviation.

### Detection of P. agathidicida *oospores in soil samples*

To determine if the identified acyl chains could serve as biomarkers for identifying *P. agathidicida* in soil, we first examined the fatty acid profile of five field collected soil samples. In each soil sample, we characterised the 12 fatty acids produced by *P. agathidicida*. Of the 12 *P. agathidicida* fatty acids, eight were readily detectable in varying quantities across samples (Suppl. Fig. 2). The addition of oospores to the soil revealed that no unique *P. agathidicida* fatty acids were detectable (Suppl. Fig. 3). However, the relative percent of the long-chain polyunsaturated fatty acid 20:5ω3, which was present in low quantities in soil (sup Figure 2) and is relatively abundant in *P. agathidicida* (Fig. 1), increased linearly with increasing oospores added to soil (Suppl. Fig. 3). In contrast, the percent of 20:4ω6 remained relatively stable with increasing numbers of oospore added (Suppl. Fig. 3). With this observation, we determined the molar quantity of 20:5ω3 and 20:4ω6 and examined the ratios of these fatty acids in each sample. The ratio of 20:5ω3 to 20:4ω6 in soil samples alone is <1 (Table 1). In *P. agathidicida* oospores, this ratio is >4 (Table 1). Thus, the presence of *P. agathidicida* in soil should lead to an increase in the ratio of 20:5ω3 to 20:4ω6 relative to *P. agathidicida-*free soil. As oospores were added to soil, the molar quantity of 20:5ω3 and 20:4ω6 both increased (Fig. 3, Suppl. Fig. 3). However, this change was greater for 20:5ω3 than for 20:4ω6 leading to a greater ratio of 20:5ω3 to 20:4ω6 as oospores increased (Table 1). In *P. agathidicida-*free soil, the 20:5ω3 to 20:4ω6 ratio was 0.92. This ratio shifted to 1.06 at the lowest concentration of added oospores (25,000), and increased to 1.4 at the highest concentration of added oospores (200,000).

**Table 1:** Ratios of 20:5ω3 to 20:4ω6 fatty acids in soil, soil with *P. agathidicida* oospores added, and *P. agathidicida* oospores alone.

|  | Soil<br>alone | 25k<br>oospores<br>in soil | 50k<br>oospores<br>in soil | 100k<br>oospores<br>in soil | 200k<br>oospores<br>in soil | 100k<br>oospores<br>alone |
| --- | --- | --- | --- | --- | --- | --- |
| 20:5ω3/20:4ω6 | 0.92 | 1.06 | 1.12 | 1.41 | 1.4 | 4.05 |

The ratios were determined by dividing the average concentration of 20:5ω3 by 20:4ω6 for each condition across five biological replicates.

**Figure 3:**
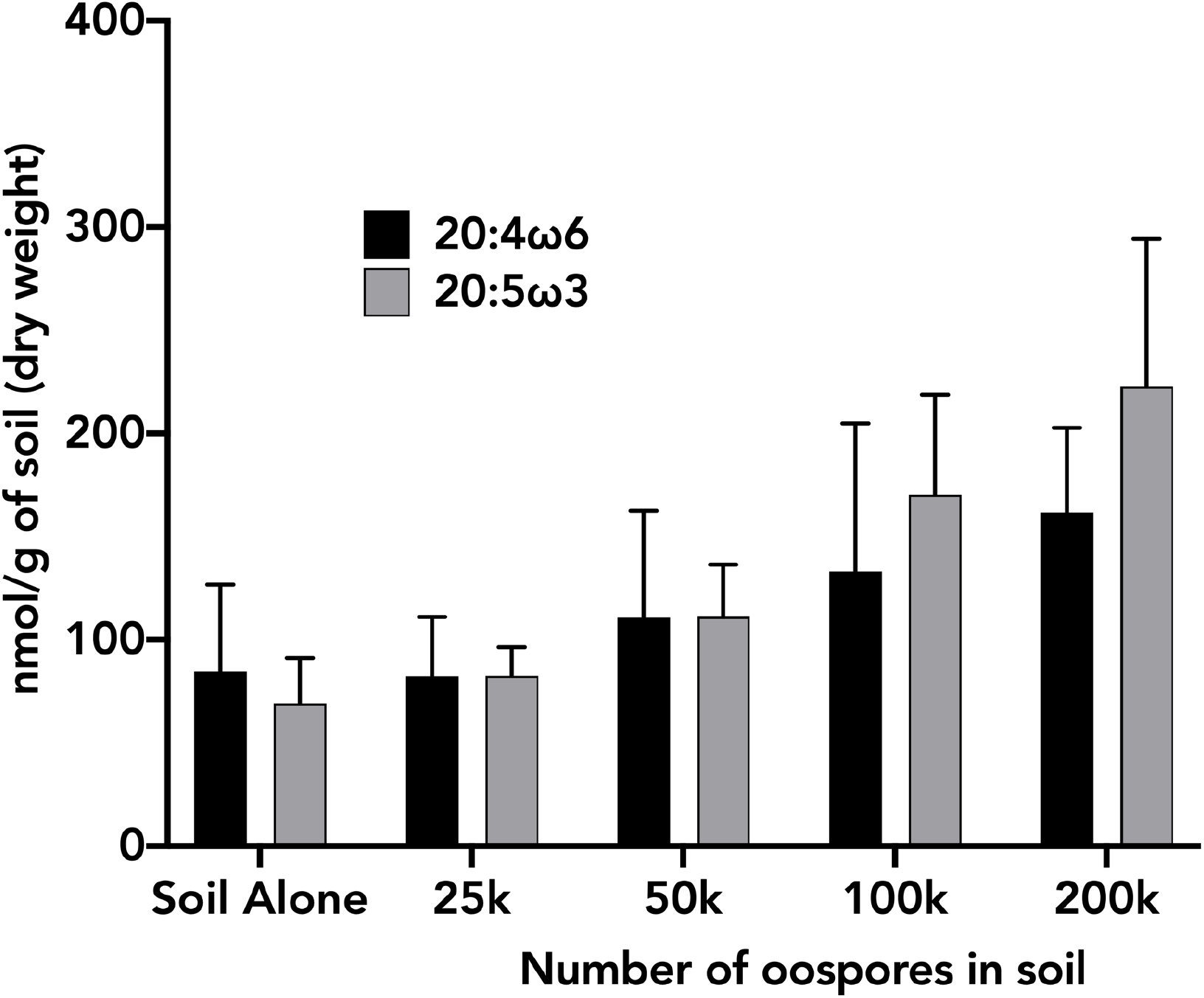
Quantification of 20:4ω6 and 20:5ω3 fatty acids in soil samples with *P. agathidicida* oospores added. Oospores were added at varying concentrations to 0.5 g of soil containing a 19:0 fatty acid internal standard. FAMEs were then produced and analysed from each sample, and the concentration of 20:4ω6 and 20:5ω3 was determined as nmol/0.5 g of soil. The values are an average of five biological replicates. Error bars indicate standard deviation.

## Discussion

Rapid and reliable diagnostics are essential when trying to manage diseases in both humans and plants. In the case of kauri dieback, tracking the spread of the *P. agathidicida* is particularly difficult due to the latent onset of disease symptoms (Bradshaw *et al*., 2020). In this study, we present the first FAME profile of *P. agathidicida*. We also assessed the potential of FAME analysis as a tool for detecting *P. agathidicida* in soil. A significant advantage of a FAME approach relative to soil baiting and PCR-based methods, is the potential to quantify the pathogen in the sample. Overall, the FAME profile of *P. agathidicida* is largely consistent with the FAME profiles of other *Phytophthora* species. The five most abundant *P. agathidicida* fatty acids (14:0, 16:0, 18:2ω6, 18:1ω9, and 20:5ω3) are also the five most abundant fatty acids in six other species of *Phytophthora* (Larkin & Groves, 2003, Duan et al., 2013). The similarity of *P. agathidicida* and other studied *Phytophthora* species is particularly interesting as *P. agathidicida* falls into the recently categorized (but as of yet understudied) *Phytophthora* clade five (Weir *et al*., 2015); our data suggest that lipid profiles may be conserved across different clades of *Phytophthora*.

While we identified no unique FAME biomarker for *P. agathidicida*, we show that the relative quantities 20:4ω6 and 20:5ω3 in soil samples can be used as an index of the likelihood that *Phytophthora* species are present. Our data reveal that the addition of increasing amounts of *P. agathidicida* oospores to soil leads to an increase in the ratio of 20:5ω3 to 20:4ω6. Thus, the ratio of 20:5ω3 to 20:4ω6 can function as a biomarker indicating the presence or absence of *Phytophthora* species in forest soils. This builds on FAME analysis performed on *P. sojae* where long-chain polyunsaturated fatty acids were detected above background soil levels when zoospores were added to soil (Yousef et al., 2012). Overall these findings suggest that ratios of long-chain polyunsaturated fatty acids can function as biomarkers for detecting *Phytophthora* on a genus level. Detecting *Phytophthora* as a genus is useful and can potentially be used in conjunction with soil bating assays or other molecular diagnostics for species confirmation. For example, large scale soil sampling and screening of samples via FAME analysis could be used to identify samples with 20:5ω3: 20:4ω6 > 1. These samples could subsequently be subjected to the soil-baiting assay to confirm the presence of *P. agathidicida*. This is similar to the “funnel and filter” model used for screening soils contaminated with *P. ramorum* (Smart et al., 2021). In that example, soils were pre-screened for the presence of *Phytophthora* using an immunosorbent assay. Soils positive for *Phytophthora* were then examined using qPCR to determine if *P. ramorum* was present.

The work presented here highlights the potential of the ratio of 20:5ω3 to 20:4ω6 to function as a tool for the detection of *Phytophthora* in soils. Nevertheless, further studies and analyses could help to optimize this promising diagnostic tool. Our results suggest that basic qualitative FAME analysis is not sufficient for differentiating *P. agathidicida* from other *Phytophthora* species. However, previous studies indicate that FAME analysis can be used to distinguish not only *Phytophthora* species but isolates within a species as well (Larkin & Groves, 2003, Duan *et al*., 2013). For example, cluster analysis based on FAME profiles was used to differentiate cultured *P. cactorum, P. citrophthora, P. cinnamomi, P. cryptogea*, and *P. nicotianae* (Duan et al., 2013). In another study, using similar techniques, individual isolates of cultured *P. infestans* were differentiated (Larkin & Groves, 2003). This suggests that a more detailed characterisation of *P. agathidicida* FAMEs may provide sufficient information for distinguishing *P. agathidicida* from other *Phytophthora* species. Additionally, utilising extraction techniques that target specific groups of lipids such as phospholipids or neutral lipids may help to further enrich target fatty acids such as 20:5ω3 and reduce background signals from complex environmental samples like soil (Drenovsky *et al*., 2004).

In contrast to FAME analysis, several DNA based diagnostic tools have been explored for use in detecting *P. agathidicida* and other *Phytophthora* species. These include both conventional qPCR and loop-mediated isothermal amplification (LAMP) methods (Than *et al*., 2013, McDougal *et al*., 2014, Hansen *et al*., 2016, Winkworth *et al*., 2020). Recently, Winkworth *et al*. combined aspects of the existing *P. agathidicida* soil-baiting assay with LAMP to reduce the overall sample processing time (Winkworth *et al*., 2020). LAMP is a DNA amplification-based method that uses primers to amplify regions of the genome-specific to an organism of interest (Wong *et al*., 2018). Efficient amplification can lead to a detectable signal, such as color-change, that can often be assessed quickly and on-site spectrophotometrically. LAMP assays efficiently detected *P. agathidicida* immediately following baiting (Winkworth *et al*., 2020). This variation on standard soil-baiting effectively eliminated the final week of the assay, shortening the process to two weeks. It is unclear whether or not LAMP would be useful at directly testing soil samples for the presence of *P. agathidicida*.

Currently, our knowledge of the geographical spread of *P. agathidicida* is limited primarily to the presence of visible disease symptoms combined with soil-baiting assays (Bradshaw *et al*., 2020). More thorough knowledge of the full range of *P. agathidicida* could potentially help answer questions regarding disease resistance in kauri trees and environmental factors that lead to infection. The knowledge gap is mainly due to the lack of a quick, cost-effective, and simple diagnostic tool. While effective, soil-baiting is not amenable to high-throughput, widespread testing. Both FAME analysis and DNA based molecular diagnostics are promising but have limitations. Further development of these techniques in conjunction with soil-baiting would help to expand our knowledge of the distribution of *P. agathidicida* in New Zealand enabling better management of kauri dieback disease.

## Acknowledgements

We’d like to thank Natascha Lewe for her assistance with fatty acid extractions and Mike Fairhurst for his assistance in isolating oospores.

