## Supplemental Figures for "The Fatty Acid Methyl Ester (FAME) profile of *Phytophthora agathidicida* and its potential use as diagnostic tool"

**Affiliations:**
[1]School of Biological Sciences, Victoria University of Wellington, New Zealand
[2]Centre for Biodiscovery, Victoria University of Wellington, New Zealand
[3]School of Chemical and Physical Sciences, Victoria University of Wellington, New Zealand
[4]Centre for Biodiversity and Restoration Ecology, Victoria University of Wellington, New Zealand

***Corresponding author**: Address: School of Biological Sciences, Level 2, Te Toki a Rata Building, Victoria University of Wellington, Wellington, 6012; Tel: +64 2102468945;

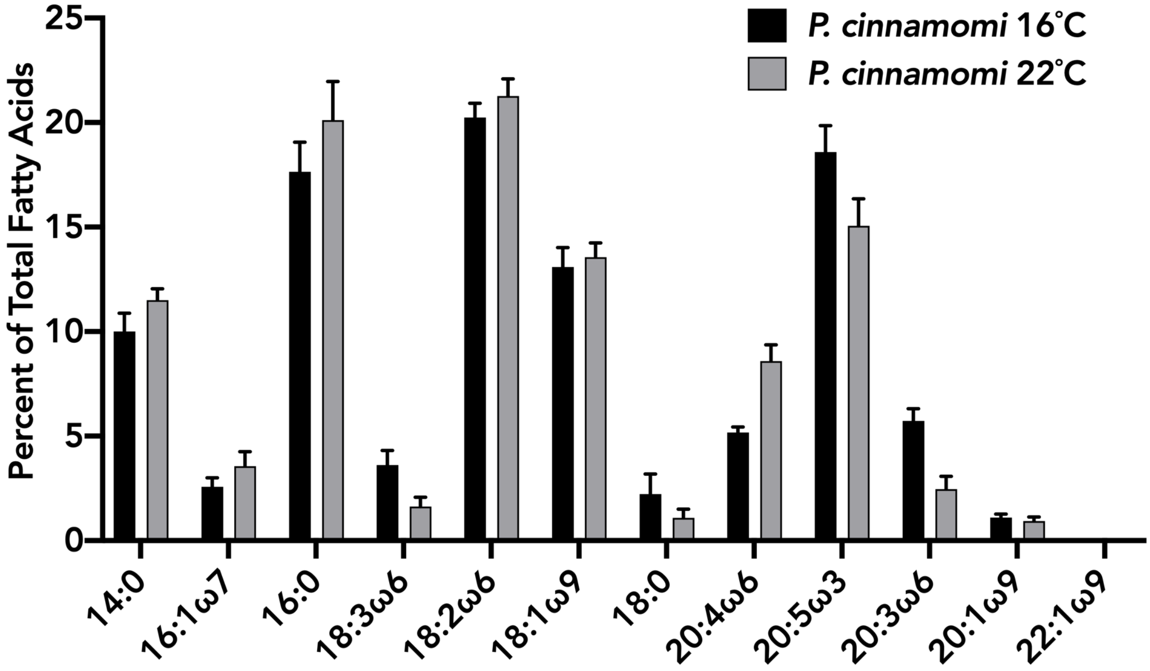


**Suppl. Figure 1: Fatty acid profile of *P. cinnamomi* at varying temperatures.** FAMEs were produced and analysed from mycelia of *P. agathidicida* grown at 16 °C and 22 °C. The percent of each fatty acid is the average relative percent of five biological replicates. Error bars indicate standard deviation.

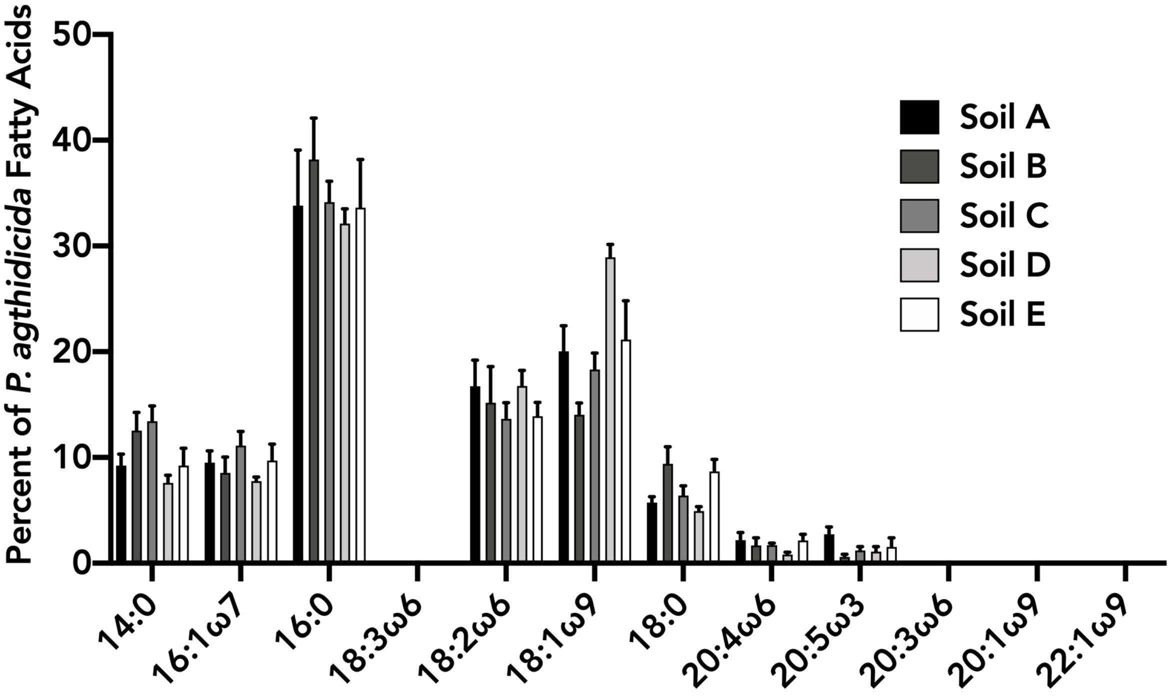


**Suppl. Figure 2: Relative percentages of *P. agathidicida* fatty acids in five soils.** Soil from five different locations in Wellington, New Zealand was collected, freeze-dried and stored at -20 °C. FAMEs were then produced and isolated from 0.5 g of each sample. Each of the *P. agathidicida* fatty acids identified in Fig. 1 were identified and quantified as a relative percentage. The values are an average of five biological replicates. Error bars indicate standard deviation.

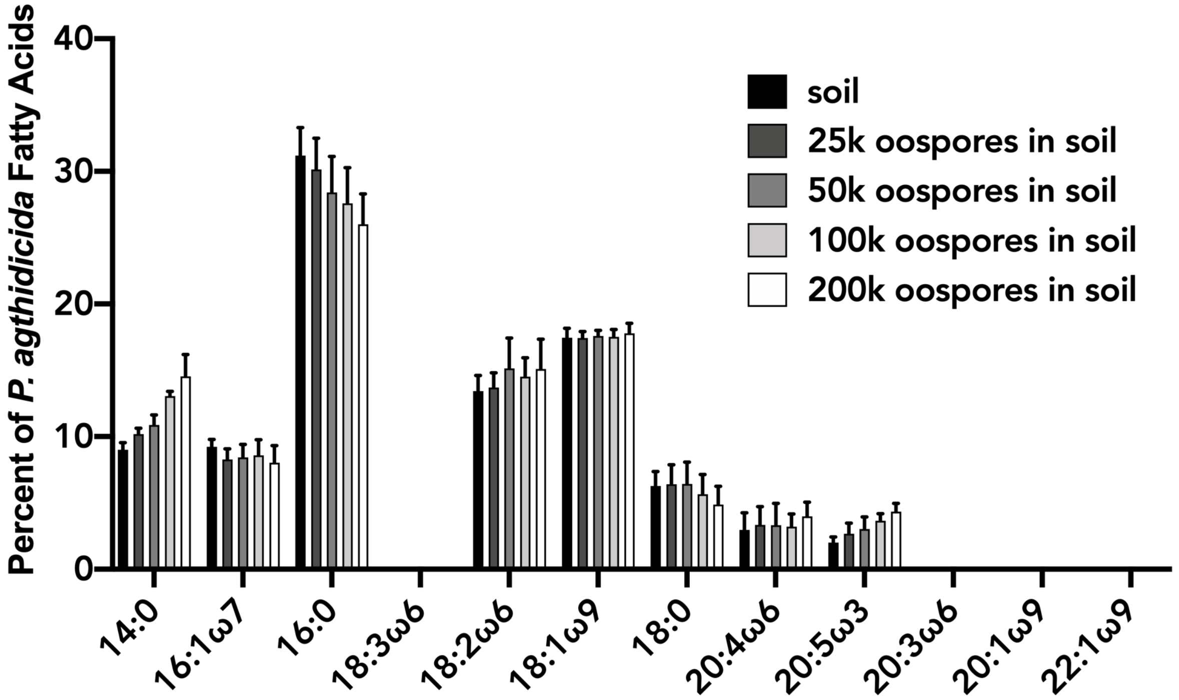


**Suppl. Figure 3: Relative percentages of *P. agathidicida* fatty acids in soil alone and soil amended with varying concentrations of oospores.** Oospores were added at varying concentrations to 0.5 g of soil containing a 19:0 fatty acid internal standard. FAMEs were then produced and analysed from each sample, and the relative percent of each of the 12 *P. agathidicida* fatty acids was determined. The percentages are an average of five biological replicates. Error bars indicate standard deviation.
